# Iterative Multiscale Molecular Dynamics: Accelerating Conformational Sampling of Biomolecular Systems by Iterating All-Atom and Coarse-Grained Molecular Dynamics Simulations

**DOI:** 10.1101/2025.02.16.638568

**Authors:** Hung N. Do, S. Gnanakaran

## Abstract

All-atom molecular dynamics (AA MD) simulations remain the gold standard for simulating biomolecular dynamics at an atomistic level for decades. However, they often cannot attain the time scales required to sufficiently sample the many biological processes of interest, especially those that involve large biomolecular systems with high energy barriers. Here, we combine the strengths of all-atom (AA) and coarse-grained (CG) MD simulations to develop the iterative multiscale MD (iMMD) simulation workflow in OpenMM by iterating between AA and CG MD simulations to enhance the sampling of biomolecular conformations. The AA-CG-AA iterations are repeated over multiple cycles, facilitating the accelerated sampling of biomolecular conformations at a CG level, while the finer atomistic interactions are refined with AA simulators. We demonstrate the enhanced sampling ability of iMMD over multiple AA-CG-AA iterations on four representative systems, including two globular proteins (fast-folding variant of Trpcage and Z-matrix protein of Mammarenavirus lassaense (LASV)) and two membrane proteins (Torpedo nicotinic acetylcholine receptor (nAChR) and REGN7663 Fab binding to the CXCR4 receptors in a heterogeneous cholesterol/phosphatidylcholine membrane lipid bilayer). In particular, iMMD captures the folding of the fast-folding Trpcage and Z-protein of LASV starting from their extended conformations as well as the binding of the REGN7663 Fab binding to the CXCR4s and dimerization of the CXCR4 receptors within the heterogenous membrane, within few AA-CG-AA iterations. The development of iMMD should allow for unprecedented simulations of biomolecules at much longer biological timescales to explore the biological processes of interest.

## Introduction

All-atom molecular dynamics (AA MD) has served as a powerful computational technique for exploring biomolecular dynamics at an atomistic level for decades^1^. However, conventional MD (cMD) simulations are computationally intensive, and despite the advances in computing hardware^2, 3^, many biological processes of interest cannot be sufficiently sampled in reasonable amount of time due to their large conformational landscapes and high energy barriers^4^.

Several strategies have been developed over the last several decades to overcome the challenges facing cMD^5^. One of such strategies involves the development of enhanced sampling methods that allow for the smoothening of the free energy landscapes and accelerate the transitions from one conformational state to another^6-11^. Another strategy simplifies the representations of biomolecules to facilitate the sampling of the conformational landscape. In particular, coarse-grained (CG) models represent biomolecules based on beads^12^, which are approximating groups of atoms. By reducing the dimensionality in MD simulations, CG simulations can be performed at elevated simulation speeds and much longer timescales compared to the AA simulations, at the expense of the atomistic details of the simulations^12^. Seemingly, the AA and CG simulations have their own strengths and weaknesses, and if combined correctly, we can retain the atomistic details provided by the AA simulations and employ the accelerated sampling capability provided the CG simulations.

In this work, we combined the strengths of the AA and CG simulations to develop the iterative multiscale molecular dynamics (iMMD) workflow by iterating between AA and CG MD simulations. The AA-CG-AA iterations were repeated over multiple cycles, facilitating the accelerated sampling of biomolecular conformations at a CG level, while the finer atomistic interactions were refined with AA simulators. iMMD was developed to be highly automatic, applicable towards a wide range of biomolecular systems rather than specific systems, and accessible to a large group of users. The enhanced sampling ability of iMMD was demonstrated on four model biomolecular systems, including two globular proteins (i.e., the fast-folding variant of Trpcage and the Z-matrix protein of Mammarenavirus lassaense (LASV)) and two membrane proteins (i.e., the Torpedo nicotinic acetylcholine receptor (nAChR) and the CXCR4s in complex with the REGN7663 Fabs in the heterogenous cholesterol/phosphatidylcholine (CHOL/POPC) lipid bilayers).

## Materials and Methods

### Workflow of the iterative multiscale molecular dynamics (iMMD)

The current implementation of iMMD is in the OpenMM simulation package^13, 14^ but can be easily adopted for GROMACS^15^. The initial input for iMMD can be a simple atomistic PDB structure of the protein of interest if the simulation system is mere solution or only involves common membrane phospholipid molecules, including DLPC, DLPE, DMPC, DOPC, DPPC, POPC, and POPE (**Figure 1**). iMMD in OpenMM can then automatically embed the protein in the phospholipid membrane of choice and/or solvate the simulation system with water and ions. If the simulation system involves non-phospholipid molecules, the user should set up simulation systems outside iMMD and provide the topology and coordinate files to start iMMD simulations.

**Figure 1.**
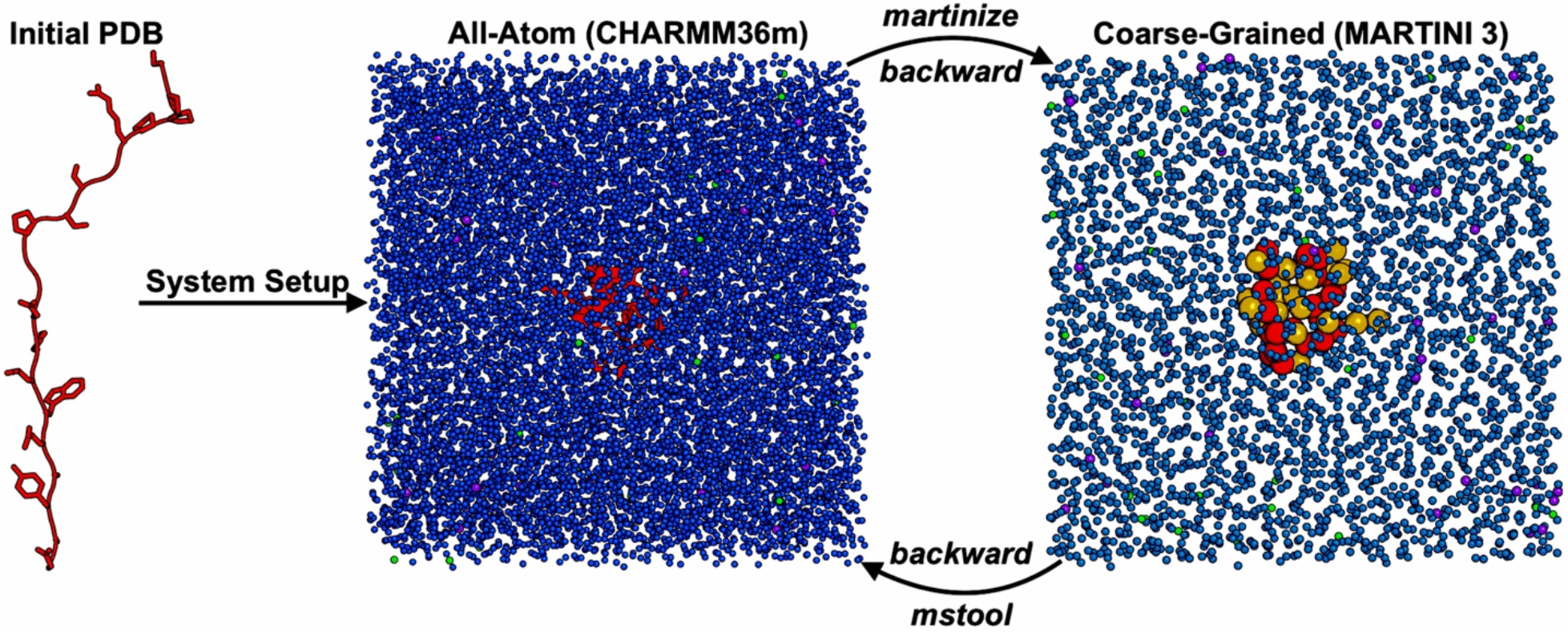
Scheme of the iterative multiscale molecular dynamics (iMMD) workflow. Given an initial PDB structure, iMMD automatically sets up the simulation system in aqueous solution or simple membrane lipid bilayer. The all-atom (AA) MD simulation is then carried out, using the CHARMM36m as the default force field parameter set. The final frame of the AA simulation is converted to its coarse-grained (CG) representation using the *Martinize2* framework for protein and *backward python* script for the other components of the simulation system. The CG MD simulation is then performed to facilitate the transitions from one conformational state to another, with MARTINI3 as the default force field parameter set. The final frame of the CG simulation is converted back to its AA representation using the *mstool python* package for the protein, common lipids, water, and ions and *backward python* script for the uncommon lipid molecules. The AA-CG-AA cycles are repeated to continue the iMMD simulation.

The AA simulation involves an initial short energetic minimization, followed by equilibration with the constant number, volume, temperature (NVT) and constant number, volume, pressure, temperature (NPT) ensembles, and finally a production simulation with NPT ensemble (**Figure 1**). The CHARMM36m force field^16^ is used by default in the AA simulation. The default settings for the AA simulation use periodic boundary conditions, with bonds containing hydrogen atoms restrained using the SHAKE^17^ algorithm. In addition, iMMD uses the Particle-Mesh Ewald summation^18^ with a cutoff of 9.0 Å for long-range interactions to calculate the electrostatic interactions. The Monte Carlo barostat^19^, with the default pressure of 1 atm and temperature of 310 K, is used for solution simulation systems, while the Monte Carlo membrane barostat^19^, with the default pressure of 1 atm, surface tension of 0 atm.nm, and temperature of 310 K, is used for simulation systems involving membranes. Furthermore, the X and Y directions of the membrane are set to scale by the same amount (or isotropic), whereas the Z direction of the membrane is allowed to vary freely. The Langevin Middle Integrator^20^, which combines Langevin dynamics with the LFMiddle discretization^21^, is used by default in the atomistic simulations, with a temperature set at 310 K, friction coefficient of 1 ps^−1^, and timestep of 2 or 4 fs.

The final frame of the AA simulation is obtained and converted into its coarse-grained (CG) representation, using the *Martinize2* and *Vermouth* framework^22^ for the proteins and the *backward*^23^ *python* script for lipids and other molecules (**Figure 1**). The *Insane*^24^ *python* package may be used to solvate the simulation system with the box dimension kept the same as in the AA simulation. The default settings for AA to CG conversion of the proteins using *Martinize2*^22^ uses side chain corrections, neutral termini, backbone position restraints, with a position restraint force constant of 2500 kJ/(mol.nm^2^), *DSSP*^25^ to determine the secondary structures, and the protein elastic network^26^, with an elastic bond force constant of 1000 kJ/(mol.nm^2^) and elastic bond lower and upper cutoffs of 0 Å and 9 Å. The CG simulation is performed following the same protocol as the AA simulation: starting with an energetic minimization, followed by equilibration with NVT and NPT ensembles, and finally a production simulation with NPT ensemble (**Figure 1**). The MARTINI 3 force field^27^ is used by default for the CG simulation. The default settings for the CG simulation treated the long-range interactions with reaction field using a cutoff of 11 Å and a relative permittivity of 15^28^. The settings for baro stat are identical to those from the AA simulations^19^. The Langevin Middle Integrator^20^ is also used for the CG simulations, with a default temperature of 310 K, collision frequency of 10 ps^−1^, and timestep of 20 fs.

The final frame of the CG simulation is then converted to its AA representation, using the *mstool*^*29*^ *python* package to backmap the protein and common lipid molecules, including DOPC, POPC, DPPC, DMPC, DSPC, DOPE, POPE, DOPS, POPS, DOPG, POPG, DOPA, POPA, SAPI, PI2A, PI2B, PI3A, PI3B, PI3C, and cholesterol (CHOL), water, and ions (**Figure 1**). The *backward python* script^23^ may be used to backmap uncommon lipid types and other molecules (**Figure 1**). The next AA simulation is performed, and the AA-CG-AA cycles are repeated, facilitating the accelerated sampling of biomolecular conformations at a CG level, while the finer atomistic interactions are refined with AA simulators.

### Simulation system set up and analysis

We demonstrated the acceleration capability of iMMD on four representative model systems, including two globular proteins in aqueous environment (i.e., the fast-folding variant of Trpcage and Z-matrix protein of the LASV) and two membrane bound proteins (i.e., the Torpedo nAChR in a heterogenous CHOL/POPC lipid bilayer, and binding of two REGN7663 Fabs to dimerizing CXCR4s in a CHOL/POPC lipid bilayer). We prepared the simulation systems of fast-folding Trpcage and LASV Z-matrix protein starting from the extended conformations of the 2JOF^30^ and 2M1S^31^ PDB structures, respectively. The solution simulation systems were automatically prepared by iMMD through the solvation of the proteins in 0.15 M NaCl solutions (**Figure 1**). Since the model membrane-bound systems of Torpedo nAChR and CXCR4s in complex with REGN7663 Fabs involved a non-phospholipid molecule (i.e., CHOL), we needed to embed the proteins in the heterogeneous lipid bilayers by the CHARMM-GUI webserver^32, 33^ prior to solvating the assembly in 0.15 M NaCl solutions in iMMD. We prepared the simulation systems of Torpedo nAChR and CXCR4s in complex with REGN7663 Fabs starting from the 4AQ9^34^ and 8U4S^35^ PDB structures, respectively. The missing regions within the 4AQ9 PDB structure were modeled using the SWISS-MODEL webserver^36^. The 8U4S^35^ PDB structure is a trimeric structure of CXCR4s in complex with REGN7663 Fabs, and we took two REGN7663 Fabs and two CXCR4 receptors from the PDB structure to prepare our simulation system. The two REGN7663 Fabs were removed from the extracellular pockets of the CXCR4 receptors and placed 20 Å away from the extracellular mouths of the CXCR4s, and the two CXCR4s were placed 20 Å away from each other. We primarily used the AmberTools^37^ along with the *MDAnalysis*^38, 39^ *python* package in analyzing the simulation trajectories obtained from the iMMD simulations. We used a definition of ≤ 9Å between Cα atoms of at least three residues apart to define contacts.

## Results

### Folding of Trpcage captured by iMMD

We demonstrated that iMMD was able to capture the folding of the fast-folding variant of Trpcage starting from its extended conformation in solution within two AA MD simulation iterations, for a total of 2 µs, and one short CG MD simulation iteration of 15 ns (**Figures 2** and **S1**). This is a seven-fold speed up compared to mere cMD of fast-folding Trpcage in explicit solvent^40^. Starting from its extended conformation in the “Unfolded” state (**Figure 2a**), Trpcage first formed a short helix from residue Y3 to L7 at ∼630 ns into the first AA MD simulation iteration in state “S1” (**Figures 2a** and **S1a**), whose Cα-atom root-mean-square deviation (RMSD) was ∼5.1 Å compared to the 2JOF^30^ PDB structure. The “S1” state was stabilized by a weak hydrogen bond between the N backbone atom of residue A4 and the O backbone atom of residue S11. Consequently, Trpcage was not able to overcome the “S1” state and stayed in that conformation for the remain of the first AA MD simulation iteration (**Figures 2a** and **S1a**). The final conformation of Trpcage in the first AA MD simulation was converted to its CG representation, and the short 15 ns CG MD simulation transitioned Trpcage from state “S1” to state “S2” by breaking the weak hydrogen between residues A4 and S11 (**Figure 2b**). In state “S2”, the long loop between residues K8 and P18 was no longer restricted by the interaction between residues A4 and S11, allowing for the fast transition into state “S3” at ∼50 ns into the second AA MD simulation iteration (**Figures 2b** and **S1b**). The Cα-atom RMSD of state “S3” compared to the 2JOF^30^ PDB structure was ∼3.0 Å, and Trpcage fully formed its major helix from residue Y3 to D9 in this state (**Figures 2b** and **S1b**). At ∼150 ns and especially ∼360 ns into the second AA MD simulation iteration, state “S3” transitioned into the “Folded” state (**Figure 2b-2c**), whose Cα-atom RMSD was ∼0.8 Å compared to the 2JOF^30^ PDB structure. As Trpcage folded, its helical proportion between residues Y3 and D9 increased from ∼0.7 at the end of the first AA MD simulation iteration to 1.0 through most of the second AA MD simulation iteration (**Figure 2c**), signifying the complete formation of the major helix. Meanwhile, the proportion of native contacts compared to the 2JOF^30^ PDB structure increased from ∼0.4 at the end of the first AA MD simulation iteration to ∼0.9 at ∼160 ns into the second AA MD simulation iteration as Trpcage folded (**Figure 2d**). Trpcage stayed in the “Folded” state for the rest of the second AA MD simulation iteration (**Figure 2b-2d**).

**Figure 2.**
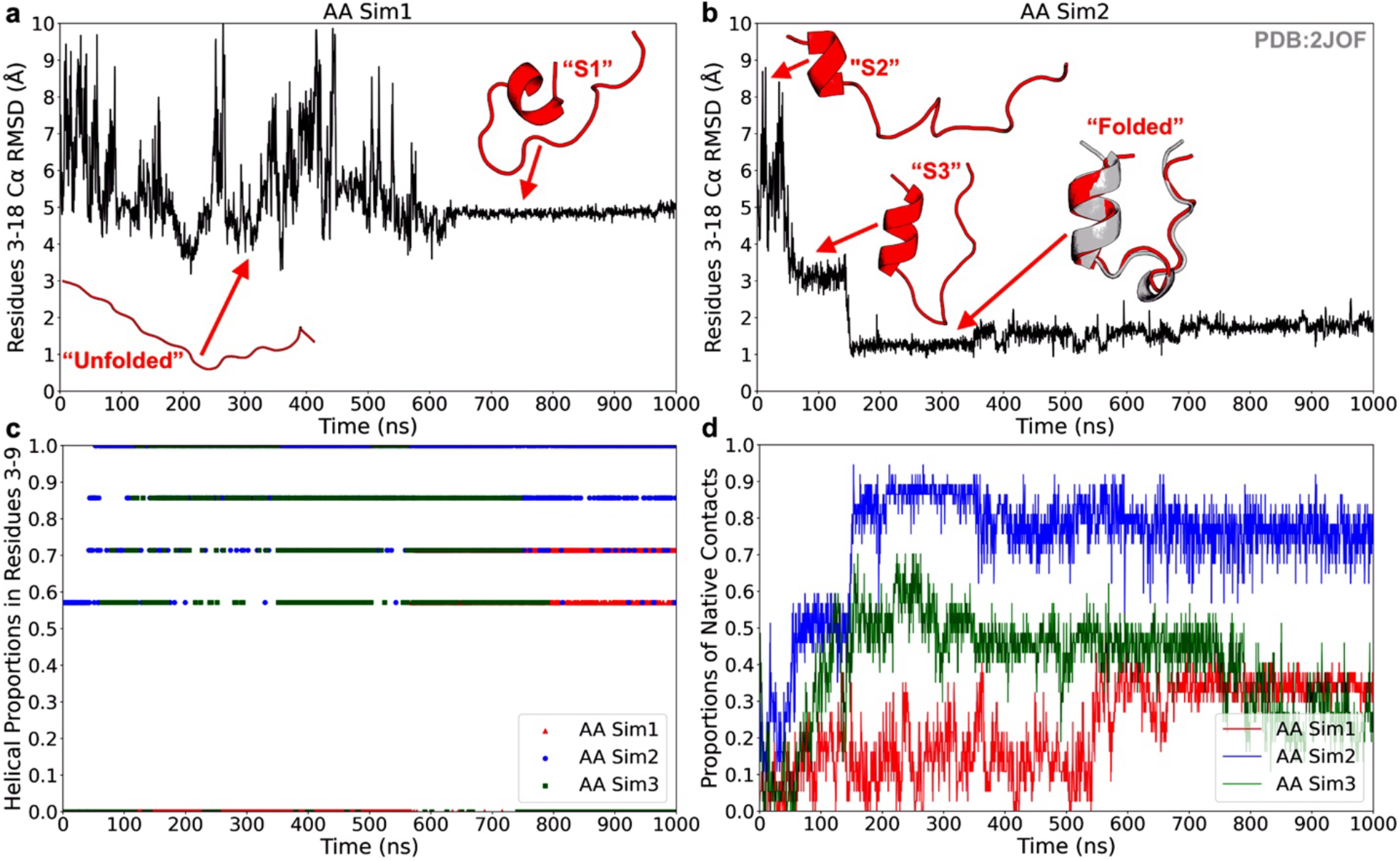
Folding of Trpcage captured by iMMD. **(a-b)** Time courses of the Cα-atom root-mean-square deviations (RMSDs) of residues 3-18 of Trpcage compared to the 2JOF PDB structure calculated from the first **(a)** and second **(b)** AA MD simulation iterations. Representative low-energy conformations obtained from the hierarchical agglomerative clustering algorithm are included in accordance with their time periods. The RMSD time courses of residues 3-9 and for the third AA MD simulation iteration are included in **Figure S1. (c)** Time courses of the helical proportions in residues 3-9 in Trpcage calculated from the first, second, and third AA MD simulation iterations. **(d)** Time courses of the proportions of native contacts in Trpcage compared to the 2JOF PDB structure calculated from the first, second, and third AA MD simulation iterations. A contact definition of ≤ 9Å between Cα atoms of at least three residues apart was used.

However, as we could not control which states our simulation system transitioned into when iterating between the AA and CG simulations, going through the second CG MD simulation iteration and back to the third AA MD simulation iteration unfolded Trpcage (**Figures 2c-2d** and **S1c-S1d**). The helical content between residues Y3 and D9 dropped to zero in the first ∼60 ns of the third AA MD simulation iteration, before fluctuating between ∼0.6 and 1.0 from the ∼80 ns to ∼800 ns mark, and finally dropped to zero in the last ∼200 ns of the iteration (**Figures 2c** and **S1d**). However, Trpcage was not able to properly refold in the third AA simulation iteration, as indicated by the high Cα-atom RMSDs of residues Y3-P18 (**Figure S1c**) and reducing proportion of native contacts compared to the 2JOF^30^ PDB structure, from ∼0.8 at the end of the second AA simulation iteration to only ∼0.3 at the end of the third AA simulation iteration (**Figure 1d**). In other words, the tertiary structure of Trpcage was not properly formed in the third AA MD simulation iteration. Eventually, the major helix between residues Y3 and D9 of Trpcage distorted at ∼760 ns into the third AA MD simulation iteration, and Trpcage returned to its “Unfolded” state (**Figure S1c-S1d**).

From our observation in the AA1-CG1-AA2 and AA2-CG2-AA3 cycles, we hypothesized that iterating between AA and CG representations likely distorted the current secondary structures of the proteins, regardless of the simulation lengths as well as position and force constraints used, due to the inconsistency between the optimal atom/bead positions in the AA and CG MD simulations. However, this distortion could be used to our advantages to transition our simulation systems out of undesirable states.

### Folding of the Z-matrix protein of the Mammarenavirus lassaense (LASV) explored by iMMD

iMMD was shown to capture the folding of the LASV Z-matrix protein, a peptide of 49 residues in size, starting from its extended conformation in solution after seven AA MD simulation iterations, for a total of 8 µs, and six CG MD simulation iterations of 100 ns each (**Figures 3** and **S2**). Due to the large protein sizes, we used longer simulation lengths for the CG simulation iterations compared to the Trpcage system to properly sample the conformational landscape. Furthermore, as the LASV Z-matrix protein primarily existed as disordered loops, except residues L51-L58 which formed a helix, we primarily paid attention to the Cα-atom RMSD of residues L51-L58 of the LASV Z-matrix protein compared to the 2M1S^31^ PDB structure, their helical contents, and their proportions of native contacts compared to the 2M1S^31^ PDB structure to determine the folding of the LASV Z-matrix protein (**Figures 3** and **S2**). During the first six AA and CG MD simulation iterations, the LASV Z-matrix protein mostly existed as disordered loops, with minor α-helices and ≤-sheets formed here and there, but no α-helix was observed between residue L51 and L58 (**Figures S2** and **3c**). The LASV Z-matrix protein started the sixth AA MD simulation in the disordered loop form of state “S1”, whose Cα-atom RMSD of residues L51-L58 compared to the 2M1S^31^ PDB structure was ∼3.8 Å (**Figure 3a**). The LASV Z-matrix protein transitioned from the “S1” to the “S2” state at ∼370 ns into the sixth AA MD simulation iteration (**Figure 3a**). In state “S2”, the Cα-atom RMSD of residues L51-L58 was ∼3.2 Å, and β-strands were formed between residues Y48-N52 and I66-P70 (**Figure 3a**). The LASV Z-matrix protein fluctuated between states “S1” and “S2” during the last ∼200 ns of the sixth AA MD simulation iteration (**Figure 3a**).

**Figure 3.**
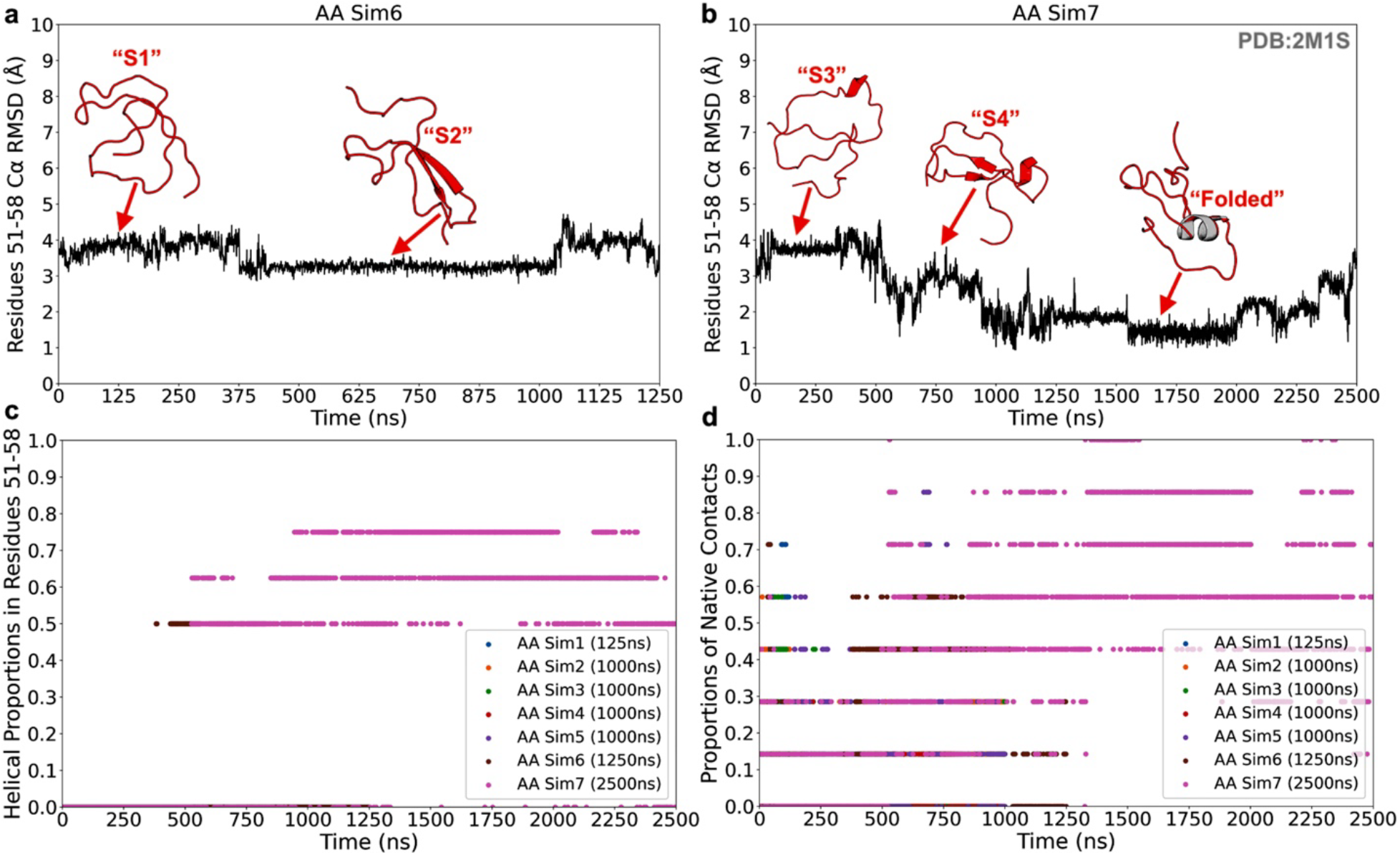
Folding of the Z-matrix protein of Mammarenavirus lassaense (LASV) uncovered by iMMD. **(a-b)** Time courses of the Cα-atom root-mean-square deviations (RMSDs) of residues 51-58 of the LASV Z-matrix protein compared to the 2M1S PDB structure calculated from the sixth **(a)** and seven **(b)** AA MD simulation iterations. Representative low-energy conformations obtained from the hierarchical agglomerative clustering algorithm are included in accordance with their time periods. The RMSD time courses calculated from the first five AA MD simulation iterations are included in **Figure S2. (c)** Time courses of the helical proportions within residues 51-58 of the LASV Z-matrix protein calculated from the seven AA MD simulation iterations. **(d)** Time courses of the proportions of native contacts within residues 51-58 of the LASV Z-matrix protein compared to the 2M1S PDB structure calculated from the seven AA MD simulation iterations. A contact definition of ≤ 9Å between Cα atoms of at least three residues apart was used.

The Z-matrix protein started the seventh and final AA MD simulation in state “S3”, whose Cα-atom RMSD of residues L51-L58 was ∼3.7 Å (**Figure 3b**). This state was primarily a disordered loop, with a minor α-helix formed between residues L56-L58 (**Figure 3b**). State “S3” of the LASV Z-matrix protein transitioned into state “S4” at ∼500 ns into the seventh AA MD simulation iteration (**Figure 3b**). The “S4” state of the LASV Z-matrix protein showed a Cα-atom RMSD of residues L51-L58 of ∼3.3 Å, and it had β-strands formed between residues C50-N52 and I66-K68 (**Figure 3b**). A minor α-helix was observed between residues T55-L58 (**Figure 3b**). As we transitioned from the “S3” to the “S4” state, the α-helix extended from residues L56-L58 to residues T55-L58. The “Folded” state of the LASV Z-matrix protein was first observed at ∼900 ns and stably at ∼1250 ns into the final AA MD simulation iteration (**Figure 3b**). In this state, the Cα-atom RMSD of residues L51-L58 of the LASV Z-matrix protein compared to the 2M1S^31^ PDB structure was ∼0.8 Å, and the α-helix extended to cover residues C53-L58 (**Figure 3b**). Residue N52 was in a turn, while residue L51 was mostly loop-like.

As the LASV Z-matrix protein gradually folded, the helical proportion between residues L51-L58 increased from 0 to ∼0.5 at ∼525 ns into the seventh AA MD simulation iteration (representing state “S4” with half of the residues from L51-L58 being helical) (**Figure 3c**). The helical content between residues L51-L58 peaked at ∼0.8 at ∼994 ns into the AA MD simulation iteration (illustrating the transition from the “S4” to the “Folded” state) and fluctuated between ∼0.5 and ∼0.8 for the remaining of the seventh AA MD simulation iteration (**Figure 3c**). In addition, iMMD was able to form most to all of the native contacts between residues L51 and L58 of the LASV Z-matrix protein compared to the 2M1S^31^ PDB structure (**Figure 3d**). In particular, the proportion of native contacts between residues L51 and L58 of the LASV Z-matrix protein compared to the 2M1S^31^ PDB structure was found to reach 1.0 at the ∼530 ns mark, between ∼1400 ns and ∼1525 ns, and between the ∼2240 ns and ∼2350 ns marks of the seventh AA MD simulation iteration (**Figure 3d**).

### Lipid equilibration in the heterogenous cholesterol/phosphatidylcholine lipid bilayer with Torpedo nAChR embedded by iMMD

CG MD simulations with MARTINI forcefield are known to facilitate lipid equilibration in heterogenous lipid bilayers compared to AA MD simulations due to their flattened free energy surfaces resulted from simplified molecular representations^12^. Furthermore, the simplified molecular representations in the CG MD simulations allow for much faster simulation speeds and longer simulation times to properly equilibrate the heterogenous lipids^12^. Here, we would like to examine the ability of iMMD to model the dynamics of membrane proteins in heterogenous lipid bilayers in terms of both protein and lipid conformations, using the Torpedo nicotinic acetylcholine receptor (nAChR) in the CHOL/POPC lipid bilayer as a model system (**Figures 4** and **S3-S6**).

**Figure 4.**
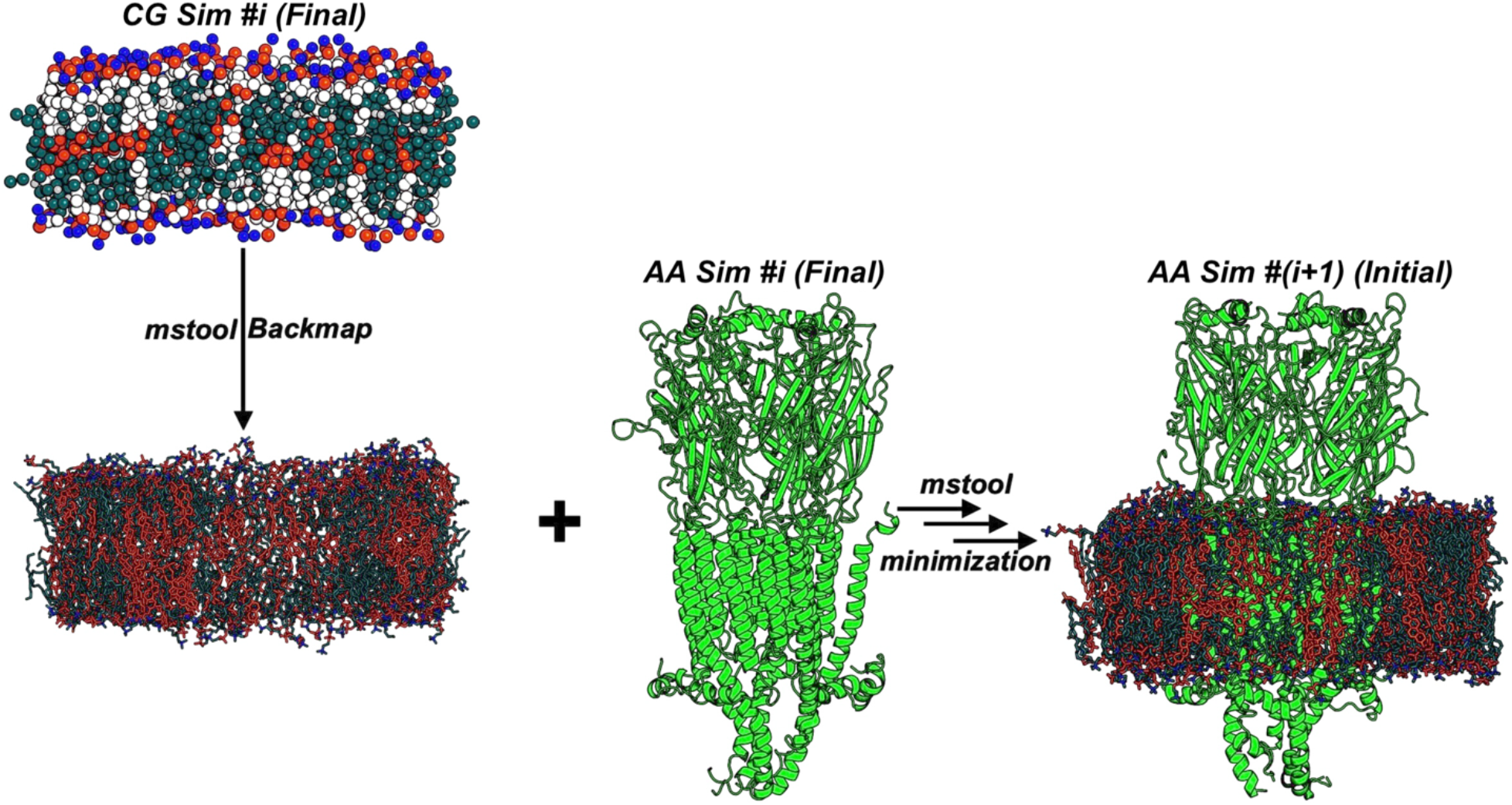
Proposed simulation protocol for iMMD simulations that involve long CG MD simulation iterations for equilibration of the heterogenous lipid bilayers. The equilibrated membrane lipid bilayers from the CG MD simulation iteration can be backmapped to their AA representations and combined with the final AA protein conformation prior to the CG MD simulation iteration using *mstool* to proceed to the next AA MD simulation iteration.

In our initial attempt, we iterated between 100 ns AA MD and 100 ns CG MD simulation iterations, with the whole simulation system (including the Torpedo nAChR and CHOL/POPC lipid molecules) cycled between their AA and CG representations using the *Martinize2* framework^22^, *backward*^23^ *python* script, and *mstool*^29^ *python* package (**Figure S3**). However, after two AA-CG-AA cycles, we observed that the secondary structures of the ligand-binding domain of the Torpedo nAChR in the solution became completely distorted (**Figure S3e**), even with the strong position and force restraints applied in the CG MD simulations. This observation confirmed our hypothesis deduced from our Trpcage simulations that iterating between AA and CG representations likely transitioned the proteins out of their current secondary structures due to the inconsistency between the force fields in the AA and CG MD simulations. The extent of this distortion might be reduced using shorter CG MD simulations. However, long CG MD simulations are required for the purpose of proper lipid equilibration in the heterogenous lipid bilayers.

Therefore, we proposed to separate the sampling of protein and lipid configurations in iMMD for systems involving heterogenous lipids (**Figure 4**). The CG MD simulations would solely be used to facilitate the equilibration of the heterogenous lipid bilayers (**Figure 4**). The protein conformations obtained from the CG MD simulations would simply be discarded. After each CG MD simulation iteration, the CG equilibrated lipid bilayers would be backmapped to their AA representations using the *mstool*^29^ and/or *backward*^23^ *python* script and combined with the final AA protein conformation prior to the CG MD simulation iteration to continue the iMMD simulation (**Figure 4**). The assembly (consisted of the AA-backmapped CG-equilibrated lipid bilayers and final AA protein conformation from the previous iteration) would be subjected to extensive energetic minimization to resolve the possible clashes between the proteins and lipids before the next AA MD simulation iteration (**Figure 4**).

Using the proposed protocol (**Figure 4**), we performed six AA MD simulations of 75-250 ns in length iterated by five CG MD simulations of 1 µs each in between on the Torpedo nAChR in the CHOL/POPC lipid bilayer. The secondary structure contents of the nAChR, including the proportions of the α-helices and β-sheets, were monitored in each AA MD simulation iteration (**Figure S4**), and we observed no difference in the proportions of the α-helices and β-sheets of the nAChR across the AA MD simulation iterations. In other words, no distortion of secondary structures took place in the nAChR as we iterated between the AA and CG simulations using the proposed protocol (**Figures S4-S5**). Using the *LiPyPhilic*^41^ python package, we calculated the lipid diffusion coefficients within the heterogeneous lipid bilayers across the first five AA and CG MD simulation iterations (**Figure S6**). The lipid diffusion coefficients were found to be higher, of 5-9 folds, during the CG MD simulations compared to the AA MD simulation iterations (**Figure S6**). In particular, the lipid diffusion coefficients during the first five AA MD simulation iterations were calculated to be ∼3.5×10^−7^ ± 5.4×10^−8^, ∼1.7×10^−7^ ± 2.2×10^−8^, ∼1.3×10^−7^ ± 1.3×10^−8^, ∼1.4×10^−7^ ± 1.4×10^−8^, and ∼1.3×10^−7^ ± 1.3×10^−8^ nm^2^/ns, respectively, while the CG lipid diEusion coeEicients were calculated to be ∼1.8×10^−6^ ± 2.5×10^−7^, ∼1.3×10^−6^ ± 1.9×10^−7^, ∼1.2×10^−6^ ± 1.7×10^−7^, ∼1.1×10^−6^ ± 2.0×10^−7^, and ∼1.1×10^−6^ ± 1.7×10^−7^ nm^2^/ns (**Figure S6**). Assuming the lipid motions during the AA and CG iterations were independent (which they were not due to the iterative nature of iMMD), the difference between the lipid diffusion coefficients between the AA and CG MD simulations were found to be statistically significant, with the p-value calculated to be less than 0.0001 using sample sizes of five (N = 5) for both AA and CG MD simulations and the unpaired student t’s test provided by GraphPad (https://www.graphpad.com/quickcalcs/ttest2/). Therefore, the CG MD simulations enhanced lipid motions within the membranes compared to the AA MD simulations. The enhanced motions were the results the flattened free energy surfaces coming from the employed MARTINI 3^27^ force field. Here, we showed the lipid molecules diffused significantly faster during CG simulations using MARTINI 3^27^ force field than AA simulations^42-44^ and justified our development of iMMD for accelerating lipid reorganization.

### Binding of the REGN7663 Fabs to dimerizing CXCR4 receptors in the heterogeneous cholesterol/phosphatidylcholine lipid bilayer explored by iMMD

In the final test of iMMD capability, we applied the method to simulate the binding of two REGN7663 Fabs to two CXCR4 receptors in the heterogeneous CHOL/POPC lipid bilayer. Modeling of protein-protein interactions remains a challenge in the field of MD^45^. Here, we aimed to test the performance of iMMD in simulating the associations of proteins in solution (i.e., REGN Fab binding to the CXCR4) and in membrane (i.e., the dimerization of the two CXCR4 receptors). Since there is currently no dimeric structure of CXCR4s in complex with REGN7663 Fabs (with 8U4R^35^ being the monomeric structure, 8U4S^35^ being the trimeric structure, and 8U4T^35^ being the tetrameric structure of CXCR4s in complex with REGN7663 Fabs), the binding conformations we found for this simulation system from iMMD might be artificial and biologically irrelevant. However, our purpose for employing this system was only to demonstrate the enhanced capability of iMMD in sampling protein-protein interactions in both solutions and membranes.

Six AA MD simulation iterations of 125-500 ns each in length iterated with five CG MD simulation iterations of 1 µs each was able to capture the binding of two REGN7663 Fabs to the extracellular domains of the two CXCR4 receptors and the dimerization of the two CXCR4 receptors (**Figures 5** and **S7-S10**). Here, we primarily used the CG MD simulations to facilitate lipid equilibration and the AA MD simulations to sample the protein-protein interactions, as described in **Figure 4**.

**Figure 5.**
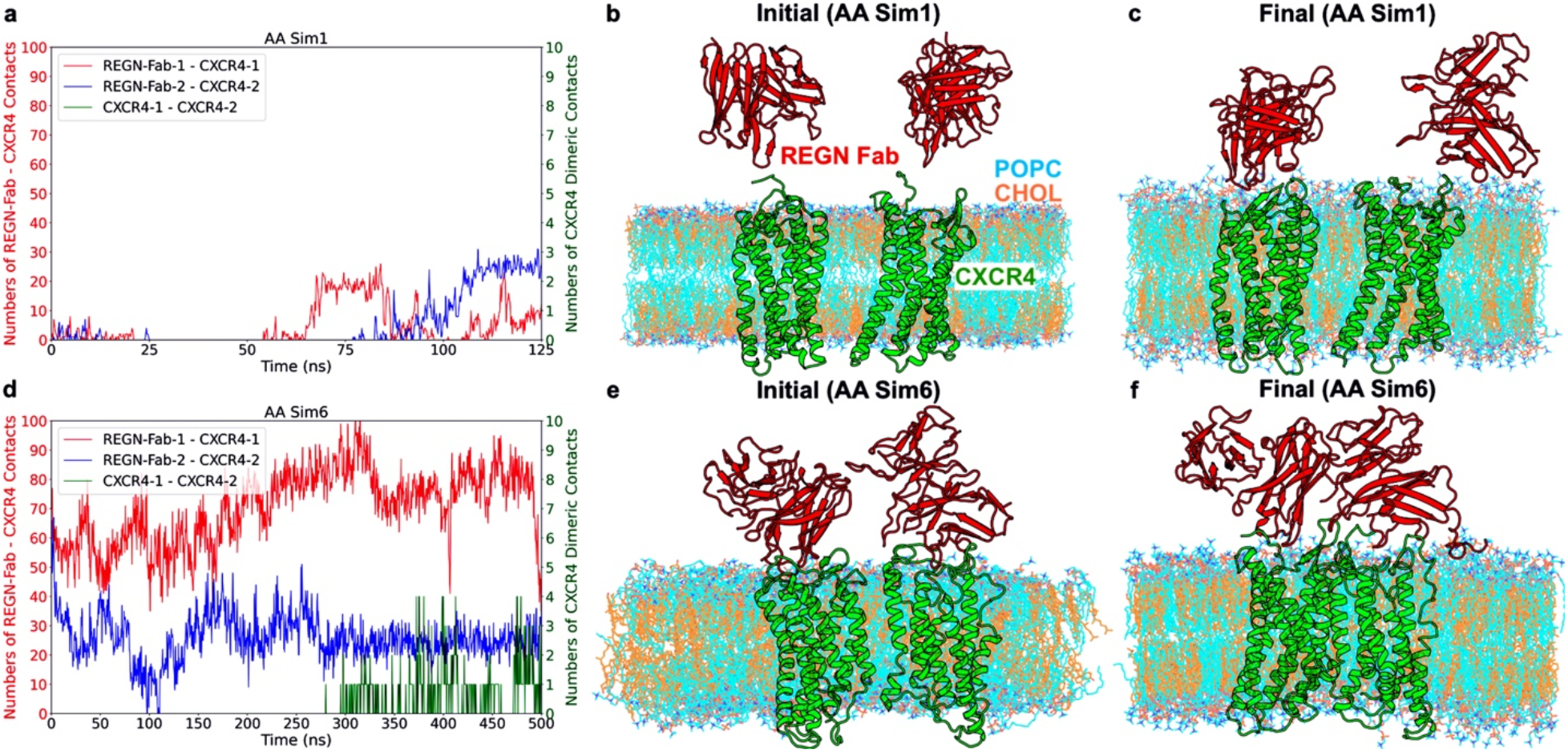
Binding of the REGN7663 Fabs to dimerizing CXCR4 receptors in the heterogenous CHOL/POPC lipid bilayer explored by iMMD. **(a)** Time courses of the numbers of contacts between the two CXCR4 receptors and between the REGN Fabs and the CXCR4s calculated from the first AA MD simulation iteration. The left y-axis is colored red and represents the number of contacts between the REGN Fabs and CXCR4s, while the y-axis is colored green and represents the number of contacts between the two CXCR4s. **(b-c)** The initial **(b)** and final **(c)** conformations of the REGN Fabs and CXCR4s in the CHOL/POPC lipid bilayer obtained from the first AA MD simulation iteration. **(d)** Time courses of the number of contacts between the two CXCR4 receptors and between the REGN Fabs and the CXCR4s calculated from the sixth AA MD simulation iteration. **(e-f)** The initial **(e)** and final **(f)** conformations of the REGN Fabs and CXCR4s in the CHOL/POPC lipid bilayer obtained from the sixth AA MD simulation iteration. A contact definition of ≤ 9Å between Cα atoms of at least three residues apart was used. The time courses of the numbers of contacts calculated from the second to fifth AA MD simulation iterations can be found in **Figure S7.** The initial and final conformations in the second-fifth AA MD simulation iterations can be found in **Figure S9**. The CXCR4s are colored green, the REGN Fabs are colored red, the POPC lipid molecules are colored cyan, and the CHOL lipid molecules are colored orange.

In the initial configuration of the first AA MD simulation iteration, the two REGN Fabs were located 20 Å away from the extracellular pockets of the CXCR4, and the CXCR4 receptors were also located 20 Å away from each other (**Figure 5b**). Towards the end of the first AA simulation iteration, we already observed from 10 to 20 contacts between the REGN-Fab-1 and CXCR4-1 as well as from 15 to 25 contacts between the REGN-Fab-2 and CXCR4-2 (**Figure 5a**), indicating the two REGN Fabs approached the CXCR4s (**Figure 5c**). However, it should be noted that few of these contacts were found in the 8U4S^35^ PDB structure, with only between ∼1% to ∼3% of the contacts being native between the REGN-Fab-1 and CXCR4-1 and none of the contacts being native between the REGN-Fab-2 and CXCR4-2 (**Figure S8**). No contact was observed between the two CXCR4 receptors throughout the first AA MD simulation iteration (**Figure 5a** and **5c**). As we transitioned into the first 1µs CG MD simulation iteration for lipid mixing and back to the second AA MD simulation iteration, we began to observe minor contact formation between the two CXCR4 receptors at the ∼53ns and ∼68ns marks (**Figures S7a**). However, it was not until the end of the final AA simulation iteration, i.e. the sixth AA MD simulation iteration, that we observed consistent contact formations between the two CXCR4 receptors (**Figures S7, S9**, and **5d-5f**). In particular, we observed from 1 to 4 contacts from the ∼300 ns mark towards the end of the AA MD simulation iteration (**Figure 5d**). At this stage, notable contacts were formed between each REGN-Fab and CXCR4 receptor, with between 35 and 102 contacts observed for REGN-Fab-1 and CXCR4-1 and up to 67 contacts observed for REGN-Fab-2 and CXCR4-2 (**Figure 5d**). However, since the simulation system was artificially constructed from the trimeric 8U4S^35^ PDB structure of CXCR4s and REGN-Fabs, most to all of the contacts were not found to be native. In particular, only up to ∼14% of the contacts between the REGN-Fab-1 and CXCR4-1 were found in the 8U4S^35^ PDB structure (**Figure S8a**), while none of the contacts between the REGN-Fab-2 and CXCR4-2 and between the two CXCR4s were found in the PDB structure (**Figure S8b**). Moreover, we also observed up to 67 nonnative contacts between the two REGN-Fabs (**Figure S10**), which could be the driving force for the CXCR4 dimerization captured in our iMMD simulation due to the simultaneous increases in number of contacts between the two REGN-Fabs and two CXCR4s observed towards the end of the sixth AA MD simulation iteration (**Figures 5d** and **S10**).

Lastly, we also calculated the lipid diffusion coefficients during the first five AA-CG MD simulation iterations of this simulation system (**Figure S11**). The lipid diffusion coefficients from the first to fifth AA MD simulation iterations were found to be ∼1.0×10^−6^ ± 5.5×10^−8^, ∼7.0×10^−7^ ± 5.5×10^−8^, ∼4.7×10^−7^ ± 3.8×10^−8^, ∼4.0×10^−7^ ± 3.7×10^−8^, and ∼3.6×10^−7^ ± 3.3×10^−8^ nm^2^/ns, respectively (**Figure S11**). The CG lipid diffusion coefficients from the first to fifth AA MD simulation iterations were calculated to be ∼3.9×10^−6^ ± 1.7×10^−7^, ∼3.2×10^−6^ ± 1.7×10^−7^, ∼3.1×10^−6^ ± 1.7×10^−7^, ∼2.9×10^−6^ ± 1.6×10^−7^, and ∼3.0×10^−6^ ± 1.7×10^−7^ nm^2^/ns (**Figure S11**). Again, if we assume that the lipid diffusion coefficients obtained from the AA and CG MD simulation iterations were independent, the p-value using the unpaired student t’s test on GraphPad (https://www.graphpad.com/quickcalcs/ttest2/) was calculated to be less than 0.0001, hence the difference between the lipid diffusion coefficients between AA and CG simulation iterations were found to be significant.

## Discussion

AA MD simulations remain the gold standard for exploring biomolecular dynamics at atomistic details. However, they are computationally expensive due to the sampling required to address the high dimensionality involved. The high energy barriers in the AA MD simulations also prevent smooth transitions from one conformational state to another. CG MD simulations allow for the accelerated sampling of biomolecular configurations due to their simplified representations of biomolecules^12^. However, they lack the atomistic details that AA MD simulations provide. CHARMM36m and MARTINI are two commonly used force fields in AA MD and CG MD simulations respectively. However, there is no relationship between these AA and CG energetic functions, so the consistency between AA and CG MD simulations cannot be guaranteed. However, each method has its own strength, and if combined correctly, we can come up with a simulation method that retains the atomistic details provided by AA simulations and the accelerated sampling capability of the CG simulations.

In this work, we combined the strengths of the AA and CG MD simulations to develop the iterative multiscale molecular dynamics (iMMD) workflow to enhance the sampling of detailed biomolecular conformations in aqueous and lipid environments. Long AA MD simulations were iterated with CG MD simulations of various simulation lengths depending on the simulation purposes over multiple cycles to allow for the accelerated transitions between conformational states during the CG simulations and refining of sampled configurations during the AA simulations (**Figure 1**). By iterating over multiple AA-CG-AA cycles rather than only one, we could achieve proper sampling of the biomolecular conformational landscapes as we facilitated multiple conformational transitions during the different CG iterations (**Figure 1**). Furthermore, since iMMD was developed using widely popular tools and force field parameter sets, the workflow should be applicable towards a wide range of biomolecular systems rather than restricted to any systems.

We demonstrated the enhanced sampling capability of iMMD over multiple AA-CG-AA iterations on four representation systems, including two globular proteins of the fast-folding variant of Trpcage and the LASV Z-matrix protein and two membrane-bound proteins of Torpedo nAChR and REGN Fab binding to the CXCR4s in the heterogenous CHOL/POPC lipid bilayers. In folding Trpcage and the LASV Z-matrix protein in solutions, we iterated long AA MD simulations with short CG MD simulations (**Figures 2-3**). The CG MD simulations were kept sufficiently short to transition the proteins from one conformational state to the next, while not to distort the secondary structures formed during the long AA MD simulations (**Figures 2-3**). iMMD was shown to capture the folding of Trpcage at seven times faster than mere cMD in solution^40^. For simulation systems that involved heterogenous lipid membranes like Torpedo nAChR and REGN Fab binding to CXCR4s, long CG simulations were required to properly equilibrate the lipid bilayers, which in turn was shown to facilitate the conformational sampling during the AA simulations. However, long CG simulations were shown to distort the current secondary structures of the proteins (**Figure S3**), even when strong position and force restraints were used. Therefore, we proposed to separate the sampling of protein and lipid configurations for the iMMD simulations of membrane proteins in heterogenous lipid bilayers (**Figure 4**). Long CG simulations were carried out to facilitate lipid equilibration only, while the protein conformations were sampled during the AA simulations (**Figure 4**). The proposed protocol was demonstrated to successfully capture the binding of two REGN Fabs to two CXCR4 receptors and the dimerization of the two CXCR4 receptors in the heterogenous CHOL/POPC lipid bilayer (**Figure 5**).

In conclusion, we developed an iterative multiscale molecular dynamics (iMMD) workflow for iterating between AA and CG MD simulations with commonly used force fields, CHARMM and MARTINI and exploiting the strengths of both. Our iMMD workflow was developed to be highly automatic for simulation systems of globular proteins and membrane proteins in simple lipid bilayers, therefore would be highly accessible to a large group of users (**Figure 1**). The workflow along with an example folder for demonstration could be found at https://github.com/lanl/iMMD. Together, the development of iMMD should allow for unprecedented simulations of biomolecules at much longer biological timescales to explore the biological processes of interest.

## Acknowledgements

This study was supported by the National Institute of Allergy and Infectious Diseases of the National Institutes of Health and by the Duke Center for HIV Structural Biology, grant number U54-AI170752-01, and Sandia National Grand Challenge Funding, CAPSIID project. The authors would also like to thank the computational resources provided by the LANL Institutional Computing. This work was performed at the Los Alamos National Laboratory, which is operated by Triad National Security, LLC, for the National Nuclear Security Administration of the U.S. Department of Energy (contract 89233218CNA000001).

