## Supporting Information for "Iterative Multiscale Molecular Dynamics: Accelerating Conformational Sampling of Biomolecular Systems by Iterating All-Atom and Coarse-Grained Molecular Dynamics Simulations"

**Figure S1.** Time courses of the Cα-atom root-mean-square deviations (RMSDs) of residues 3-9 of Trpcage compared to the 2JOF PDB structure calculated from the first **(a)** and second **(b)** all-atom (AA) molecular dynamics (MD) simulation iterations as well as residues 3-18 **(c)** and residues 3-9 **(d)** calculated from the third AA MD simulation iteration of Trpcage by iMMD.

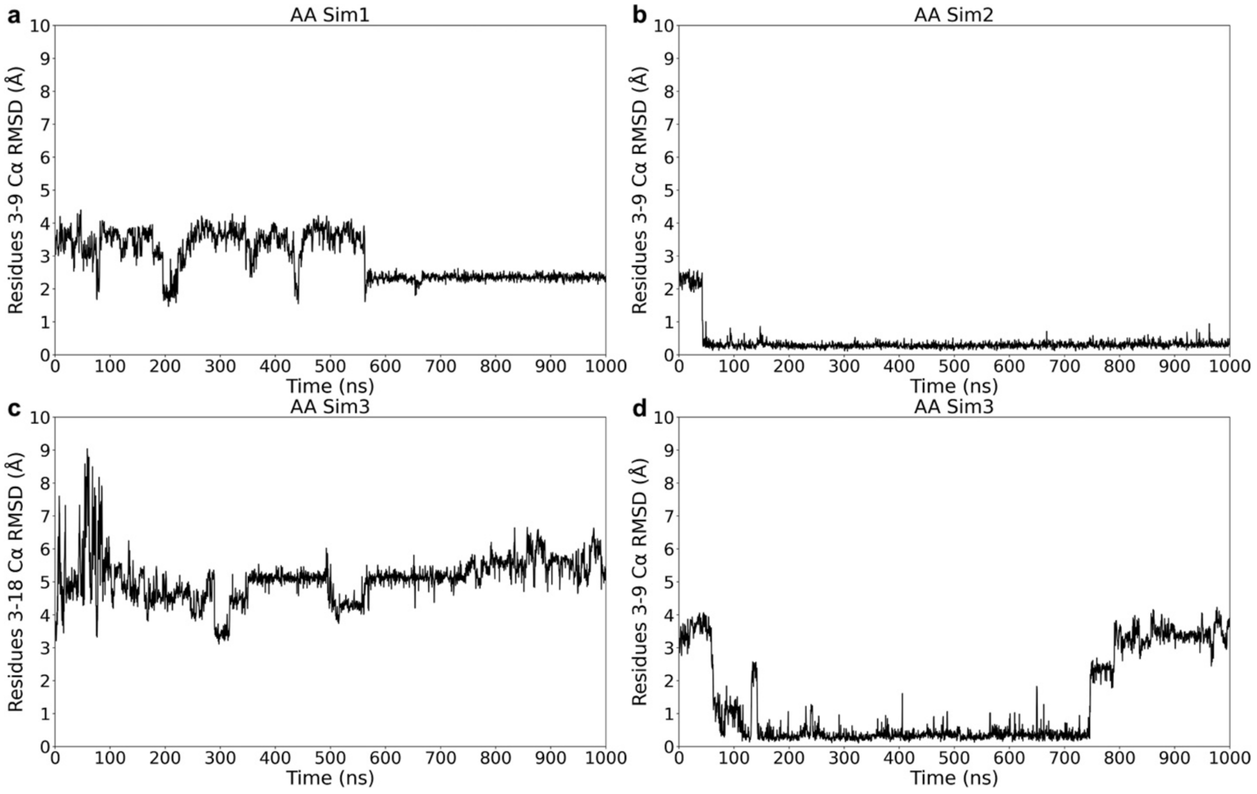

**Figure S2.** Time courses of the Cα-atom RMSDs of residues 51-58 of the Z-matrix protein of the mammarenavirus lassaense (LASV) compared to the 2M1S PDB structure calculated from the first five AA MD simulation iterations by iMMD. The initial and final conformations of the LASV Z-matrix protein in each AA MD simulation iteration are included in the left and right sides, respectively, of the corresponding panels.

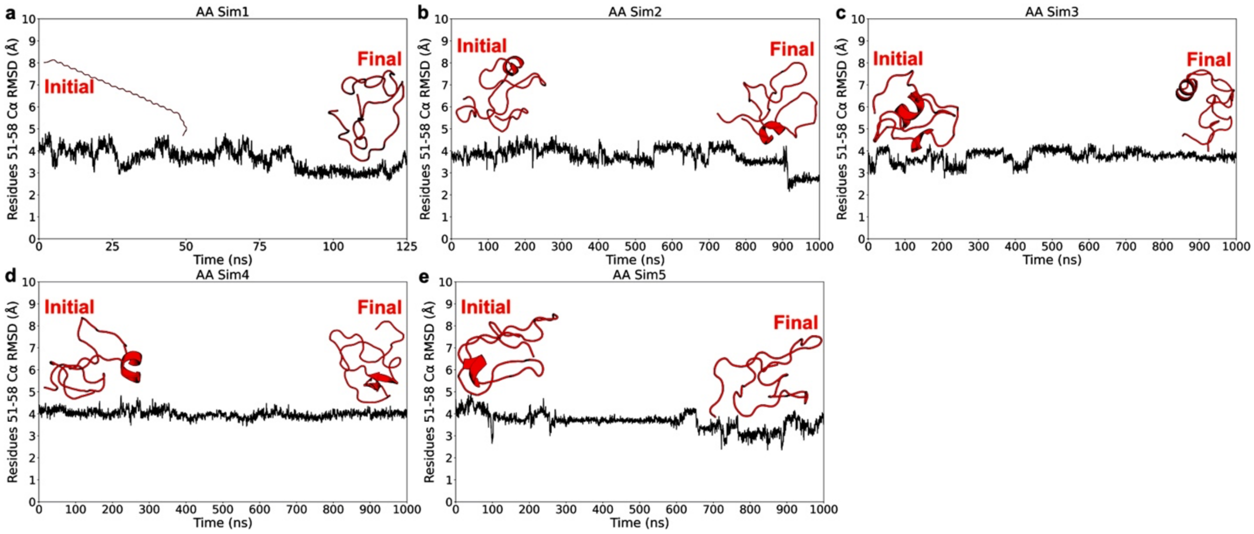

**Figure S3.** The initial and final configurations of the membrane-bound Torpedo nicotinic acetylcholine receptors (nAChRs) obtained from the AA and CG MD simulation iterations in the initial iMMD simulation of Torpedo nAChR in the heterogenous CHOL/POPC lipid bilayer, where the whole CG simulation systems (including the proteins and lipid molecules) were backmapped to their AA representations to continue the simulations. The secondary structures of the protein soluble domains became heavily distorted after few AA-CG-AA cycles. The nAChRs are colored green, the POPC lipid molecules are colored cyan, and the CHOL lipid molecules are colored orange.

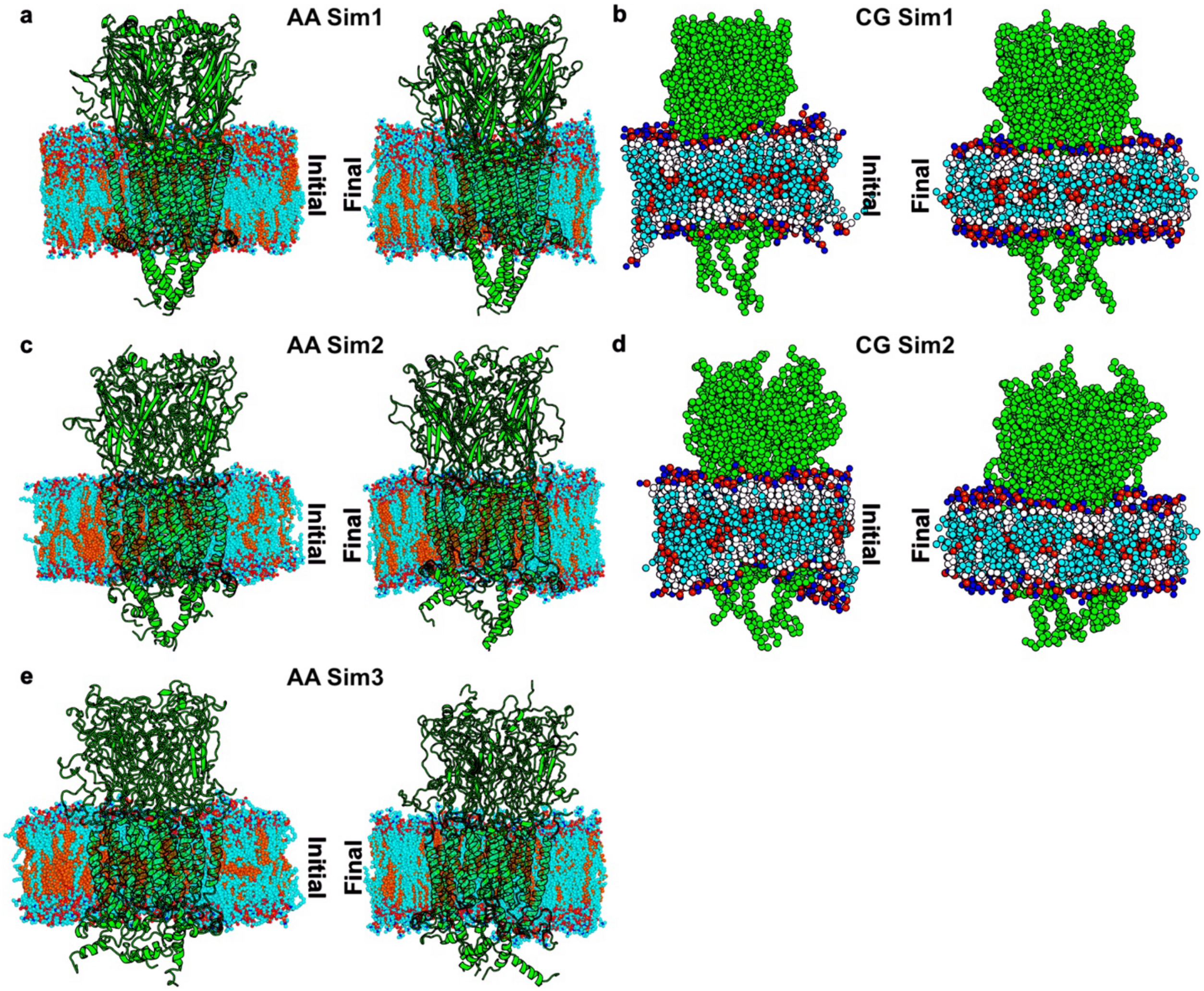

**Figure S4.** Time courses of the secondary structure proportions of the $\alpha$-helices and $\beta$-sheets calculated from the six AA MD simulation iterations in the later iMMD simulation of Torpedo nAChR in the CHOL/POPC lipid bilayer, where only the lipid molecules from the final CG simulation snapshots were backmapped and combined with the AA protein conformations from before the CG simulations to continue the simulations, following the protocol described in **Figure 4**. No secondary structure distortion was observed through the AA-CG-AA cycles in these iMMD simulations.

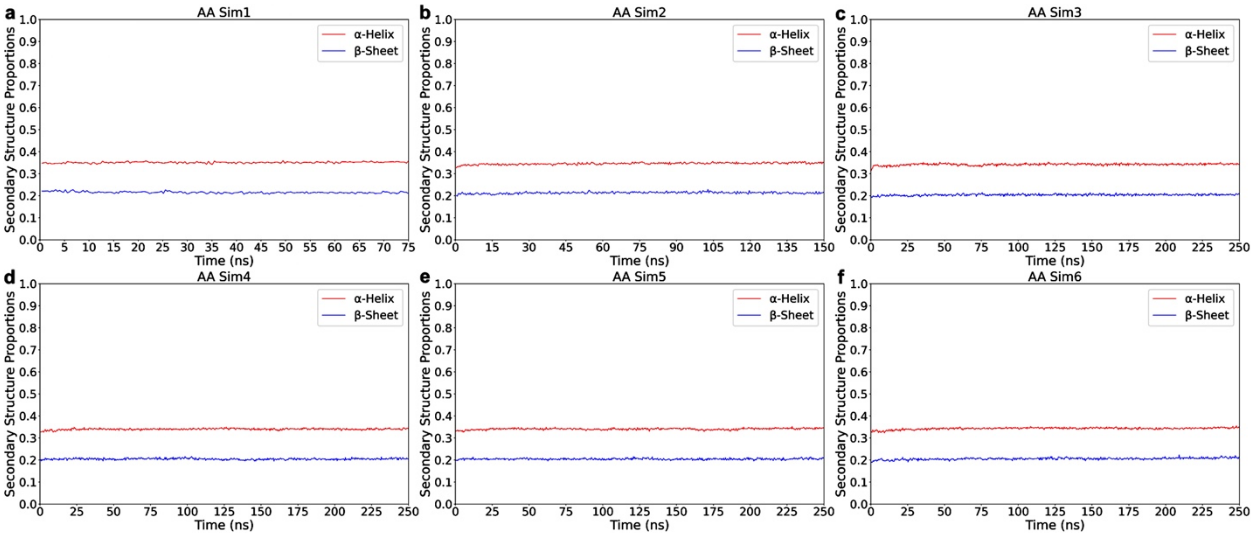

**Figure S5.** The initial and final conformations of the Torpedo nAChR in the heterogenous lipid bilayer obtained from the six AA MD simulation iterations in the later iMMD simulation. No secondary structure distortion was observed through the AA-CG-AA cycles in these iMMD simulations. The nAChRs are colored green, the POPC lipid molecules are colored cyan, and the CHOL lipid molecules are colored orange.

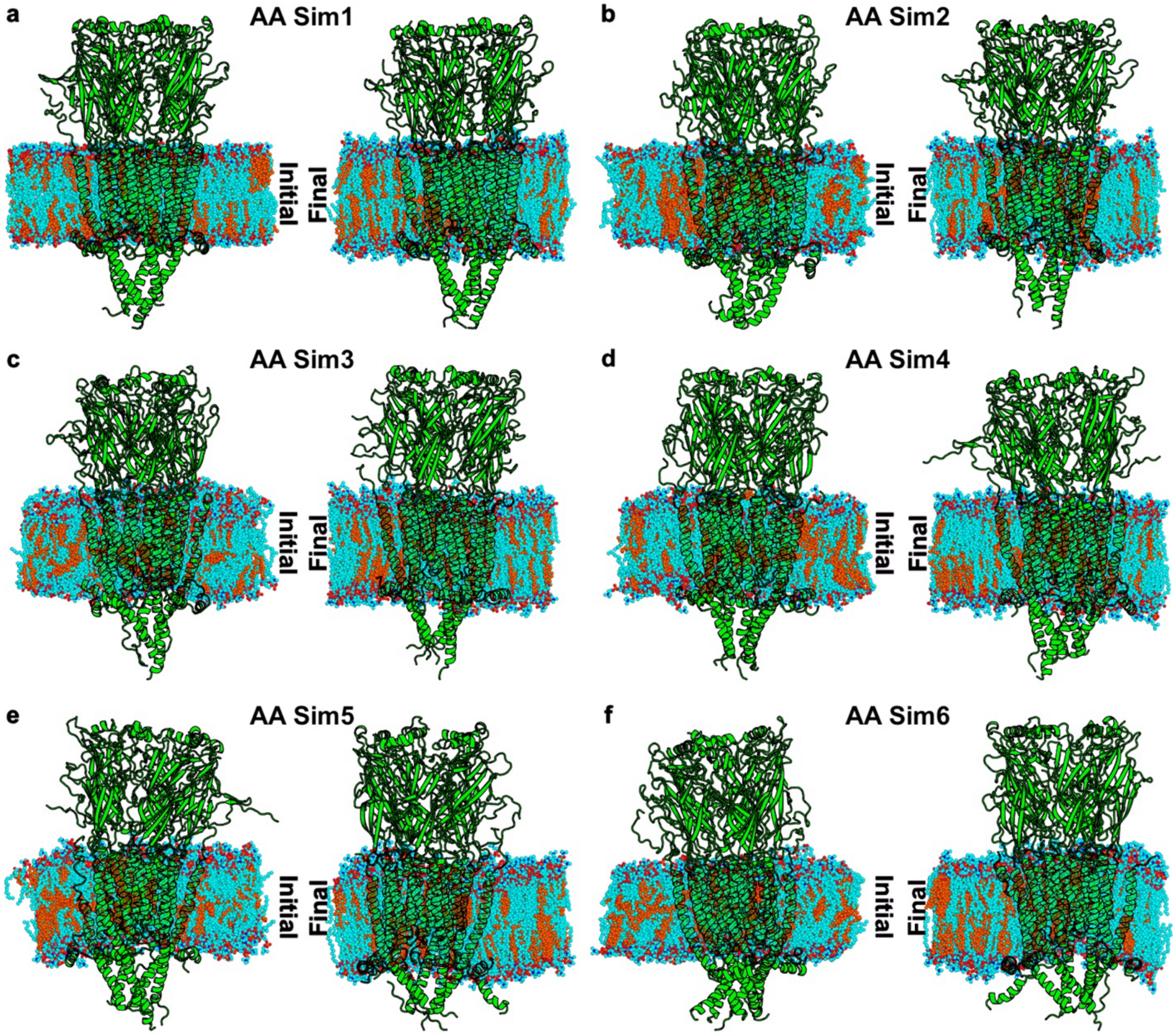

**Figure S6.** Lipid diffusion coefficients calculated from the AA and CG MD simulation iterations of the Torpedo nAChR in the heterogenous CHOL/POPC lipid bilayer by the *LiPyphilic python* package. The CG MD simulations facilitated lipid motions within the membranes compared to the AA MD simulations.

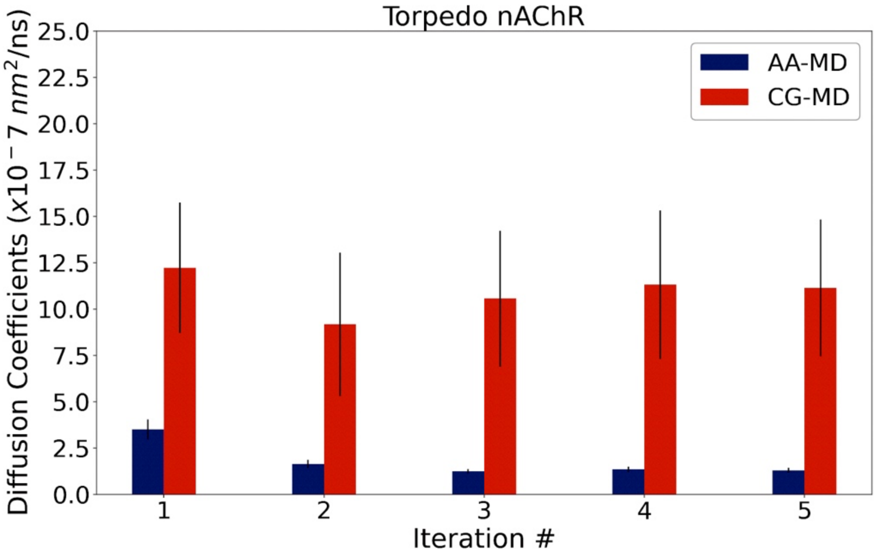

**Figure S7.** Time courses of the numbers of contacts between the REGN Fabs and the CXCR4s and between the two CXCR4 receptors calculated from the second **(a)**, third **(b)**, fourth **(c)**, and fifth **(d)** AA MD simulation iterations of the REGN Fab binding to the CXCR4s by iMMD. The left y-axis is colored red and represents the number of contacts between the REGN Fabs and CXCR4s, while the y-axis is colored green and represents the number of contacts between the two CXCR4s. A contact definition of $\leq 9Å$ between $C\alpha$ atoms of at least three residues apart was used.

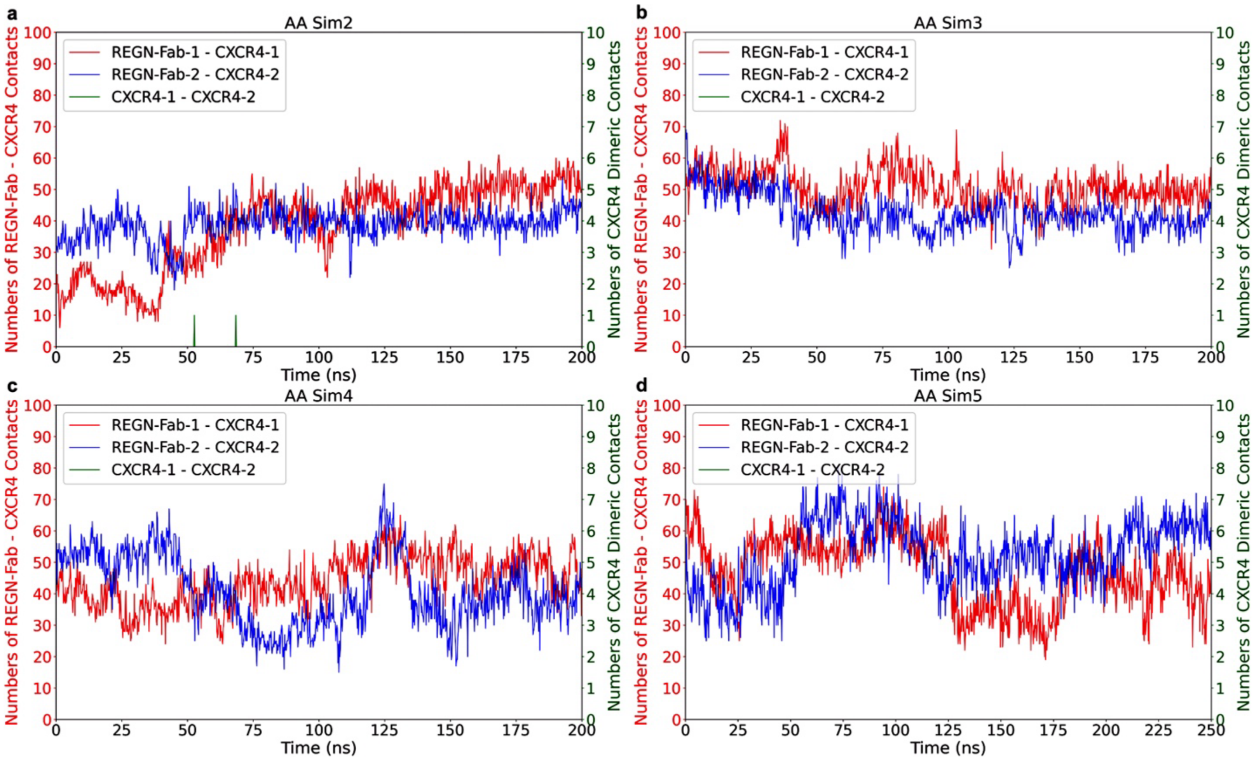

**Figure S8.** Time courses of the proportions of native contacts between the REGN Fabs and CXCR4 receptors relative to the 8U4S PDB structure calculated from the six AA MD simulation iterations of the REGN Fab binding to the CXCR4s by iMMD. A contact definition of ≤ 9Å between Cα atoms of at least three residues apart was used. It should be noted that 8U4S is a trimeric structure of CXCR4 in complex with REGN Fabs, but we only considered the dimeric complex of CXCR4 with REGN in our study.

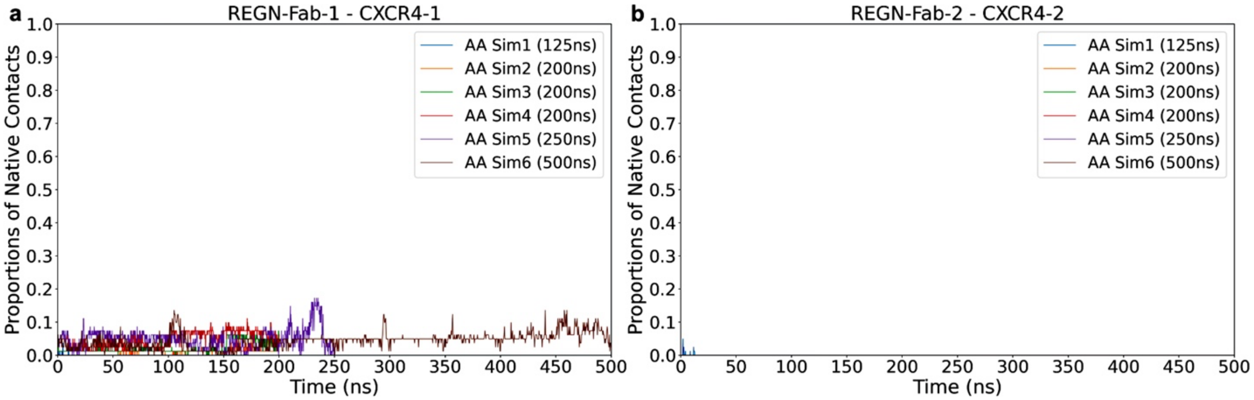

**Figure S9.** The initial and final conformations of the REGN Fabs and CXCR4s in the heterogenous CHOL/POPC lipid bilayers obtained from the second **(a)**, third **(b)**, fourth **(c)**, and fifth **(d)** AA MD simulation iterations of the REGN Fab binding to dimerizing CXCR4s by iMMD. The CXCR4s are colored green, the REGN Fabs are colored red, the POPC lipid molecules are colored cyan, and the CHOL lipid molecules are colored orange.

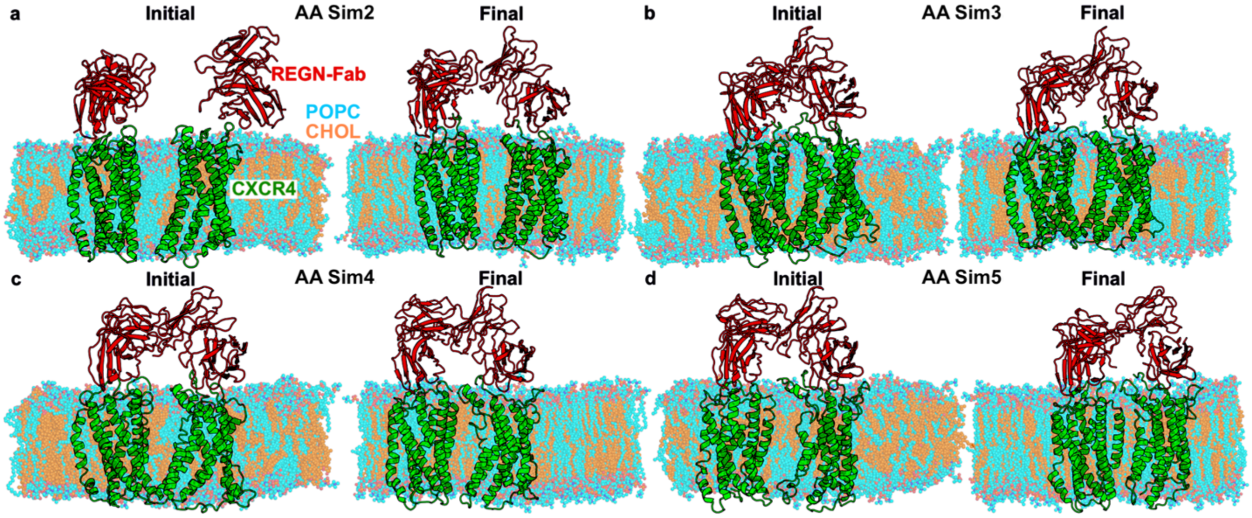

**Figure S10.** Time courses of the number of contacts between the two REGN Fabs calculated from the six AA MD simulation iterations of REGN Fab binding to the CXCR4s by iMMD. A contact definition of ≤ 9Å between Cα atoms of at least three residues apart was used.

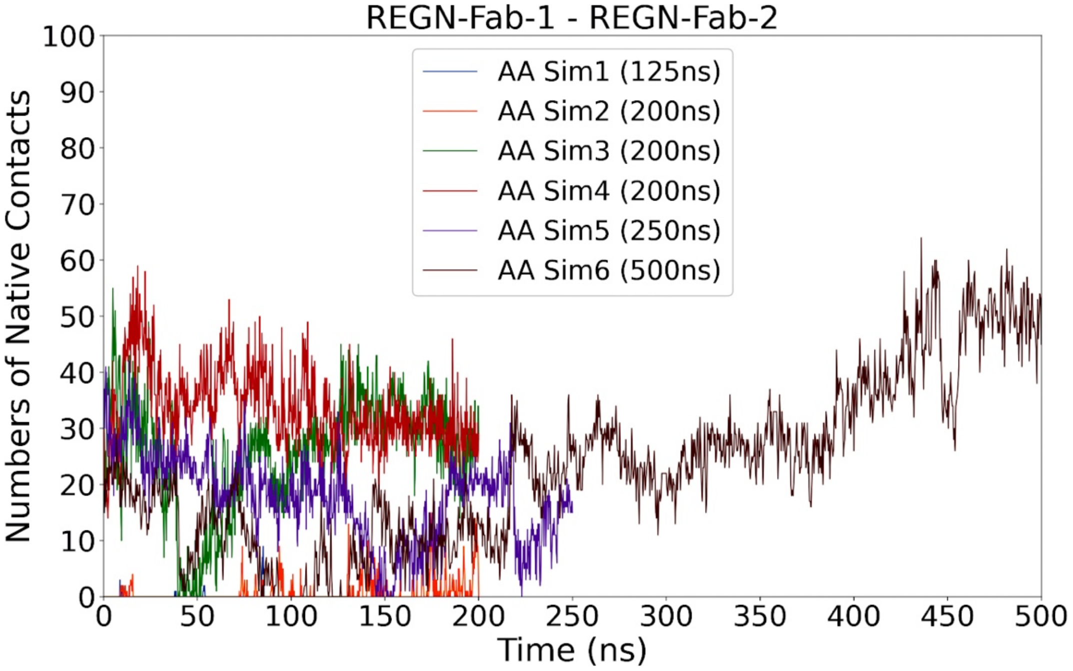

**Figure S11.** Lipid diffusion coefficients calculated from the AA and CG MD simulation iterations of the REGN Fab binding to the CXCR4s in the heterogenous CHOL/POPC lipid bilayer by the *LiPyphilic python* package. The CG MD simulations facilitated lipid motions within the membranes compared to the AA MD simulations.

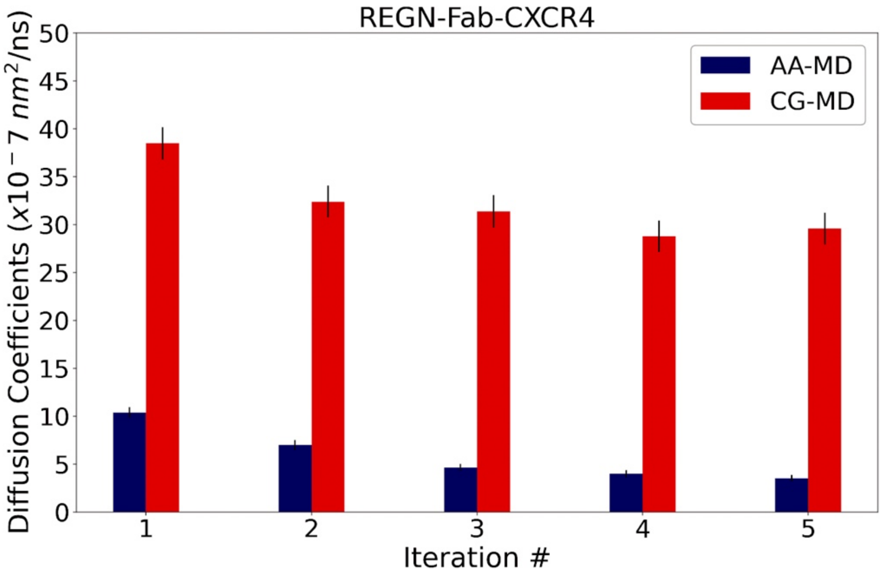
